# Neural coupling between spinal motor neurons of the first dorsal interosseous muscle during individual index finger flexion and pinch tasks

**DOI:** 10.64898/2026.04.09.717449

**Authors:** Elmira Pourreza, Hélio V. Cabral, Nijia Hu, J Greig Inglis, Mikaël Desmons, Ioannis Delis, Laura McPherson, Francesco Negro

## Abstract

**Objective:** Precision grip tasks require complex coordination of intrinsic hand muscles, yet how common synaptic inputs to motor neurons are modulated during functionally different tasks remain unclear. This study investigated whether neural coupling between motor unit spike trains in the first dorsal interosseous (FDI) muscle differs between isolated index finger flexion and precision pinch tasks.

**Approach:** Sixteen healthy participants performed isolated index finger flexion and pinch tasks at 10% and 20% of maximal voluntary contraction while high-density surface electromyography was recorded from the FDI. Motor unit spike trains were decomposed and tracked across tasks. Neural coupling was assessed using complementary methods: coherence analysis and Proportion of Common Input (PCI) index to quantify linear common oscillations in delta (1-5 Hz), alpha (5-15 Hz), and beta (15-35 Hz) frequency bands, and mutual information-based network analysis to capture nonlinear interactions.

**Main results.:** Coherence analysis and PCI revealed no significant differences between tasks across all frequency bands. In contrast, network density derived from mutual information analysis showed significantly stronger nonlinear motor unit coupling during pinch compared to isolated finger flexion (p = 0.013), independent of force level.

**Significance.:** These findings demonstrate a dissociation between linear and nonlinear measures of motor unit coupling. In particular, precision pinch tasks appear to rely on stronger higher–order common inputs and distinct neural control strategies that are not fully captured by traditional linear coherence measures. This highlights that functionally relevant precision behaviors engage additional layers of nonlinear neural coupling, offering new insight into how the nervous system adaptively modulates motor unit coordination to meet complex task demands.

## INTRODUCTION

The human hand enables a wide repertoire of grips, allowing versatile and precise interactions with the environment. The control of individual finger forces during precision grips (i.e., grips involving the thumb and one or more of the other fingers) enables holding objects between the palmar surfaces of the fingers and the opposing thumb (Napier, 1956), allowing fundamental actions in daily activities. Generating stable and controlled finger forces in such tasks requires the complex coordination of multiple intrinsic and extrinsic hand muscles (Madarshahian & Latash, 2021; Tanzarella et al., 2020, 2021; Winges et al., 2008; Santello et al., 2016). For instance, during precision grips involving the thumb and index finger (i.e., pinch), the first dorsal interosseous (FDI) and thenar muscles (abductor pollicis brevis, flexor pollicis brevis, and opponens pollicis) act as primary force generators, producing opposing force vectors that must be precisely coordinated to stabilize the thumb-index interface while modulating grip force (Cooney et al., 1985; Giurintano et al., 1995; Kozin et al., 1999; Valero-Cuevas et al., 2003). Importantly, even movements that appear isolated to a single finger rely on coordinated activation across hand muscles to stabilize joints and counteract unintended torques arising from the mechanical coupling of the digits (Madarshahian & Latash, 2021; Tanzarella et al., 2020, 2021; Cabral et al., 2025). Together, these observations highlight that both synergistic and assumed isolated finger actions depend on tightly coordinated muscle activity to achieve functional hand control.

Previous research suggests that finger force control during precision grip is achieved through modular organization of descending corticospinal inputs, rather than through independent control of each individual degree of freedom (Tanzarella et al., 2021; Gentner & Classen, 2006; Santello et al., 2016; Bizzi & Cheung, 2013). Seminal works in the hand demonstrated that lesions to the corticospinal tract (Tower, 1940; Lawrence and Kuypers, 1968; Lawrence and Hopkins, 1976) or to the primary motor cortex (Schieber and Poliakov, 1998; Liu and Rouiller, 1999) in monkeys profoundly impair the ability to perform independent finger movements, indicating that the development of precision during grasping is critically dependent on the synaptic connections in the corticospinal pathways. These findings suggest that, at the spinal level, common synaptic projections from the motor cortex to alpha motoneurons located within and across hand muscles play a critical role in coordinating grip forces. However, the degree of shared inputs appears to differ between hand muscle groups. While extrinsic hand muscles generally exhibit a higher degree of commonality, the intrinsic hand muscles express lower commonality between motoneuronal activity (Winges et al., 2008; McIsaac & Fuglevand, 2008; Tanzarella et al., 2020, 2021). This reduced synchronization, linked to the lower commonality of motoneuronal activity, is thought to support finer, low-force, independent control of individual digits, required for precision grip. Additionally, nerve block studies have demonstrated that transient pharmacological blockade of the intrinsic hand muscles’ peripheral nerves severely impairs precision grip performance (Kozin et al., 1999). Together, these results suggest that modulations in corticospinal pathways are responsible for shaping the activation patterns of intrinsic hand muscles, enabling the fine and flexible control required for precision grip tasks.

Modulations in corticospinal drive to alpha motoneurons can be estimated using time- and frequency-domain correlation techniques on motor unit spike trains during voluntary contractions (Negro & Farina, 2012). Common oscillations in motor unit discharge patterns, particularly in the delta (1-5 Hz) and beta (15-35 Hz) frequency bands, have been associated to common drive during steady contractions and corticospinal contributions, respectively (Conway et al., 1995; Baker et al., 1997; Farina & Negro, 2015; McManus et al., 2019). Therefore, estimating such common synaptic oscillations using correlation techniques provides a window to explore corticomotoneuronal contributions in muscle control. Previous studies have explored this possibility in hand control by investigating how task differences alter shared synaptic oscillations to motoneurons innervating intrinsic hand muscles. McIsaac and Fuglevand (2008) and Winges et al. (2008) demonstrated that the adductor pollicis and FDI exhibit markedly lower motor-unit synchrony during precision grip than extrinsic muscles, suggesting more selective descending inputs. In contrast, Del Vecchio et al. (2019) reported that motoneurons from the FDI and thenar muscles still received substantial common synaptic input during a non-synergistic digit task, indicating that shared descending drive can emerge between intrinsic muscles under specific task conditions. These findings suggest that intrinsic hand muscles are coordinated by shared synaptic input that is flexible and task-specific, allowing the nervous system to adapt to different grasping demands.

However, previous studies were constrained by several limitations. Some employed pairwise correlation methods that are biased by motoneuron discharge statistics (de la Rocha et al., 2007; Negro & Farina, 2012) and can only capture second-order interactions, failing to detect higher-order and nonlinear dependencies that may be critical for understanding motor unit coordination during complex tasks. Moreover, studies of intrinsic hand muscles examined limited motor unit samples or narrow force ranges and lacked direct task comparisons between isolated and synergistic actions at matched force levels. Therefore, it remains unclear whether neural coupling patterns observed during isolated finger movements are fundamentally reorganized during functionally relevant precision grip tasks, or whether task-specific differences simply reflect adjustments within the same underlying control system. To address these limitations, we analyzed shared synaptic inputs to the FDI alpha motoneurons during isolated index-finger flexion and precision pinch at matched relative force levels. We quantified common input by combining linear second-order methods (coherence and Proportion of Common Input) with higher-order nonlinear analyses (network-information framework), while tracking the same motor units across tasks. This integrated approach allowed us to characterize how common synaptic inputs are modulated as task demands change. Based on the greater coordination requirements of precision pinch and prior evidence of task-dependent modulation in extrinsic hand muscles (Tanzarella et al., 2021; Cabral et al., 2025), we hypothesized stronger common inputs during pinch.

## METHODS

### Participants

Sixteen healthy right-handed subjects (6 females; age: 29 ± 5) participated in this study. All subjects had no history of musculoskeletal or neurological disorders affecting the upper limb. Prior to data collection, participants received comprehensive information about the experimental procedures and written informed consent was obtained from all participants. The study protocol was approved by the local ethics committee (NP5890) and conducted in accordance with the latest version of the Declaration of Helsinki.

### Experimental setup and protocol

Participants were instructed to sit in front of a custom-designed pinch manipulandum (Figure 1A), with their right arm and wrist secured to a support device with the shoulder at 50°, the elbow at 150°, and the wrist in a neutral position. The manipulandum consisted of two precision load cells (SM S-TYPE-500N and 100N, Interface, U.S. & METRIC), upon which participants placed their index and thumb fingers on adjustable supports (Figure 1A). This experimental setup enabled the measurement of isometric forces exerted by the index finger and the thumb flexion. Prior to data collection, the skin overlying the first dorsal interosseous (FDI) was shaved and mildly abraded with an abrasive paste (EVERI, Spes Medica, Genova, Italy) and cleaned with water. Subsequently, a HDsEMG grid consisting of 64 electrodes (HD04MM1305; 13 rows × 5 columns; 4–mm interelectrode distance; OT Bioelettronica, Turin, Italy) was affixed over the FDI muscle belly (Figure 1A). Grid was fixed to the skin using a two-sided adhesive foam, and the electrode-skin contact was ensured by filling the foam wells with conductive paste (AC cream, Spes Medica, Genova, Italy). The reference electrode band was positioned on the right wrist.

**Figure 1:**
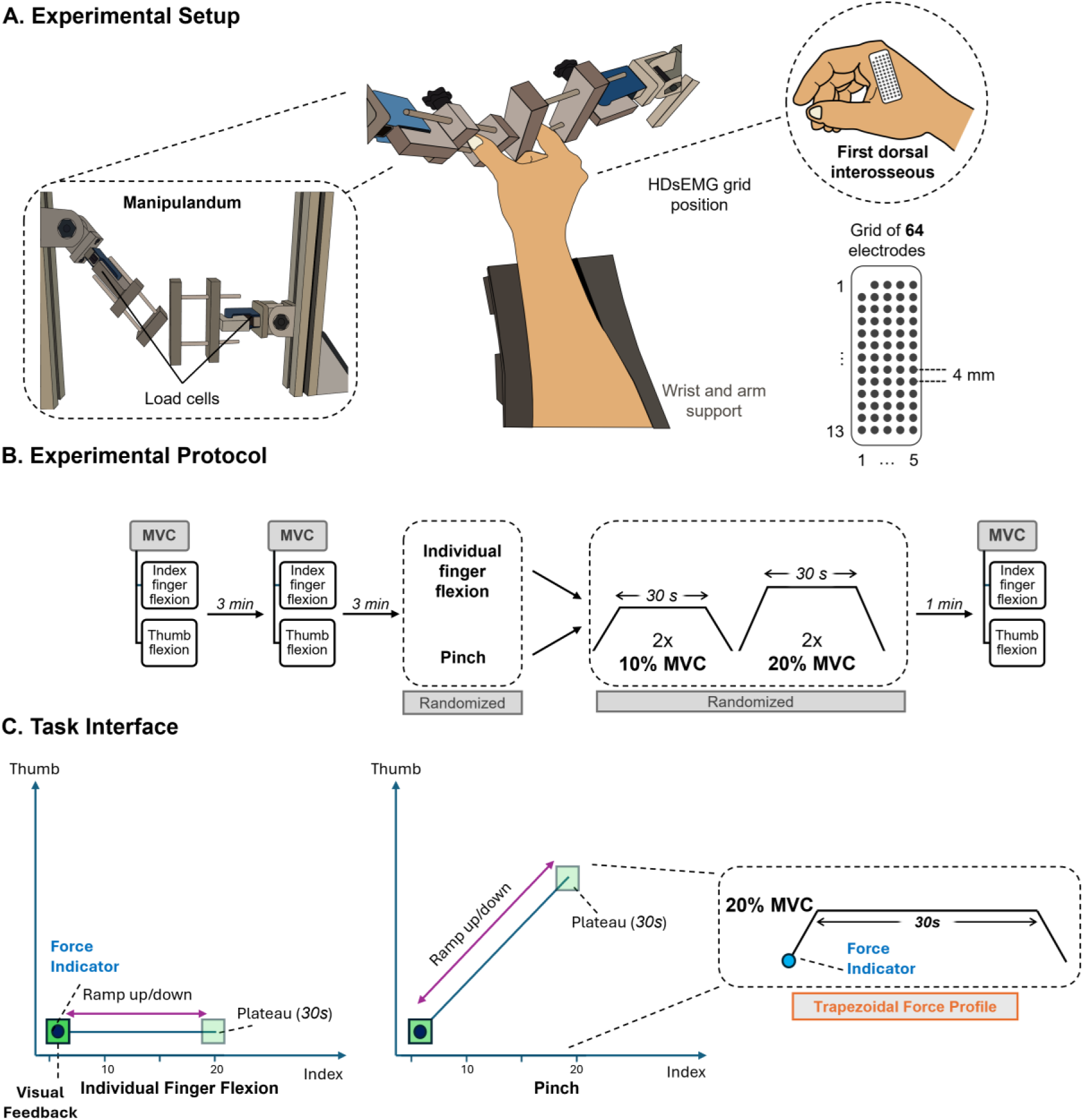
A) shows the experimental setup including a custom-made manipulandum with two load cells to record the isometric forces produced by the flexion of the index finger and thumb, the hand position during experiments, the position of HDsEMG grid on the first dorsal interosseous (FDI), and the 64-channel electrode grid arrangement. B) shows the experimental protocol where participants performed isometric contractions at 10% and 20% of maximal voluntary contraction (MVC) for individual index finger flexion and pinch tasks (randomized order). Target trajectories were visualized for the trapezoidal pattern. C) shows the task interface, where x axis represents the individual finger flexion and y axis represents the thumb movement, and the diagonal trajectory represents pinch task.

The experimental protocol is detailed in Figure 1B. Two trials of maximum isometric voluntary contractions (MVCs) for the thumb (pollex, first digit) and index finger (second digit) flexion were initially recorded separately, with a 3-min rest period between trials. The peak MVC value was used as a reference for the submaximal tasks. Subsequently, participants were asked to repeat an isometric task for two distinct conditions: (a) individual index finger flexion, (b) pinch (i.e., index finger and thumb flexion together). For each condition, they performed submaximal isometric contractions following trapezoidal profiles. Each trapezoidal profile encompassed a gradual ramp-up phase from 0% to the target force at a rate of 3% MVC/s, a plateau phase in which the target force was maintained for 30 s, and a ramp-down phase from the target force to 0% MVC at the same rate (Figure 1B). Two trials were performed for each target force level, which were set at 10% and 20% MVC. Real-time visual feedback from the exerted force was provided to assist participants in maintaining the target force levels. The order of conditions and force levels were randomized across participants. At the end of the experimental protocol, an MVC measurement was recorded again to verify that fatigue had not affected the recordings.

### Data collection

HDsEMG signals were recorded in a monopolar configuration and digitized at 2,000 Hz using a wireless amplifier (16 bits; 10-500 Hz bandwidth; Sessantaquattro+, OT Bioelettronica, Turin, Italy). Force signals from the index finger flexion and thumb flexion were amplified with a gain of 100 (low pass: 500Hz, high pass: DC coupled; Forza amplifier, OT Bioelettronica, Turin, Italy) and recorded along with the feedback signals through auxiliary channels of the HDsEMG amplifiers. Feedback was provided using a visual interface that displayed a moving target square following a trapezoidal force profile (Figure 1C). The square appeared in the positive quadrant of an x–y Cartesian coordinate system, with its trajectory originating from (0, 0). Its path was configured according to the specific motor task: for index finger flexion, the target moved along the x-axis; for thumb flexion, along the y-axis; and for pinch tasks, along the diagonal (x–y) line. Subjects were instructed to modulate their force output so that a corresponding circle on the screen closely tracked the moving target. The target square advanced in a straight line at a constant speed of 3% MVC per second. Upon reaching the specified target force, it paused for 30 seconds before returning directly to the origin to initiate the subsequent cycle. Moreover, the dimensions of the square were scaled relative to % MVC to maintain an allowable margin of error of ± 2% MVC.

### Data analysis

#### Motor unit decomposition

Prior to analysis, the root-mean-square error (RMSE) between the force profile and the feedback signal was computed for each trial and the one with lower RMSE (i.e., highest force-feedback matching) was selected for further processing. Motor unit analysis was restricted to the 30-s plateau phase. Initially, HDsEMG signals were bandpass filtered with a third-order Butterworth filter (20-500 Hz cutoff frequencies), after which low-quality channels (e.g., those containing artifacts) were visually identified and excluded from further analysis. Motor unit spike trains were then extracted from the remaining HDsEMG signals using a convolutive blind source separation algorithm (Negro et al., 2016). This technique has been extensively employed to evaluate the activity of individual motor units from hand muscles (Negro et al., 2016; Cabral et al., 2024). Following automated detection, each motor unit spike train was carefully inspected by an experienced investigator to identify and correct any missing or falsely detected discharges (Negro et al., 2009; Hug et al., 2021). Only spike trains with a silhouette value greater than 0.87, indicative of a high decomposition accuracy, were included in the subsequent analyses (Negro et al., 2016).

#### Motor unit tracking

Separately for each force level (10% and 20% MVC), motor units were tracked across the two tasks (index finger flexion and pinch) by reapplying the motor unit separation vectors, following procedures used in previous studies (Oliveira & Negro, 2021; Cabral et al., 2024). Briefly, decomposing HDsEMG signals with independent component analysis entails computing a separation matrix whose columns consist of distinct separation vectors corresponding to individual motor units (Negro et al., 2016). Each of these vectors serves as a spatiotemporal filter that, when convolved with the HDsEMG signals, provides the discharge times of the motor units. To optimize motor unit tracking, all the separation vectors derived from both conditions (individual index finger flexion and pinch) were systematically applied to the concatenated HDsEMG signals of the conditions, maximizing the number of matched motor units (Figure 2). Duplicate matches occurred because some motor units were independently decomposed in both conditions and therefore appeared twice after the matching step. Duplicates were identified as pairs with overlap in discharge times (> 30% common spikes). After identification, the unit with the lower coefficient of variation of inter–spike intervals (CoV-ISI) was retained for further analysis (Chen & Zhou 2022, Martínez-Valdés 2017).

**Figure 2:**
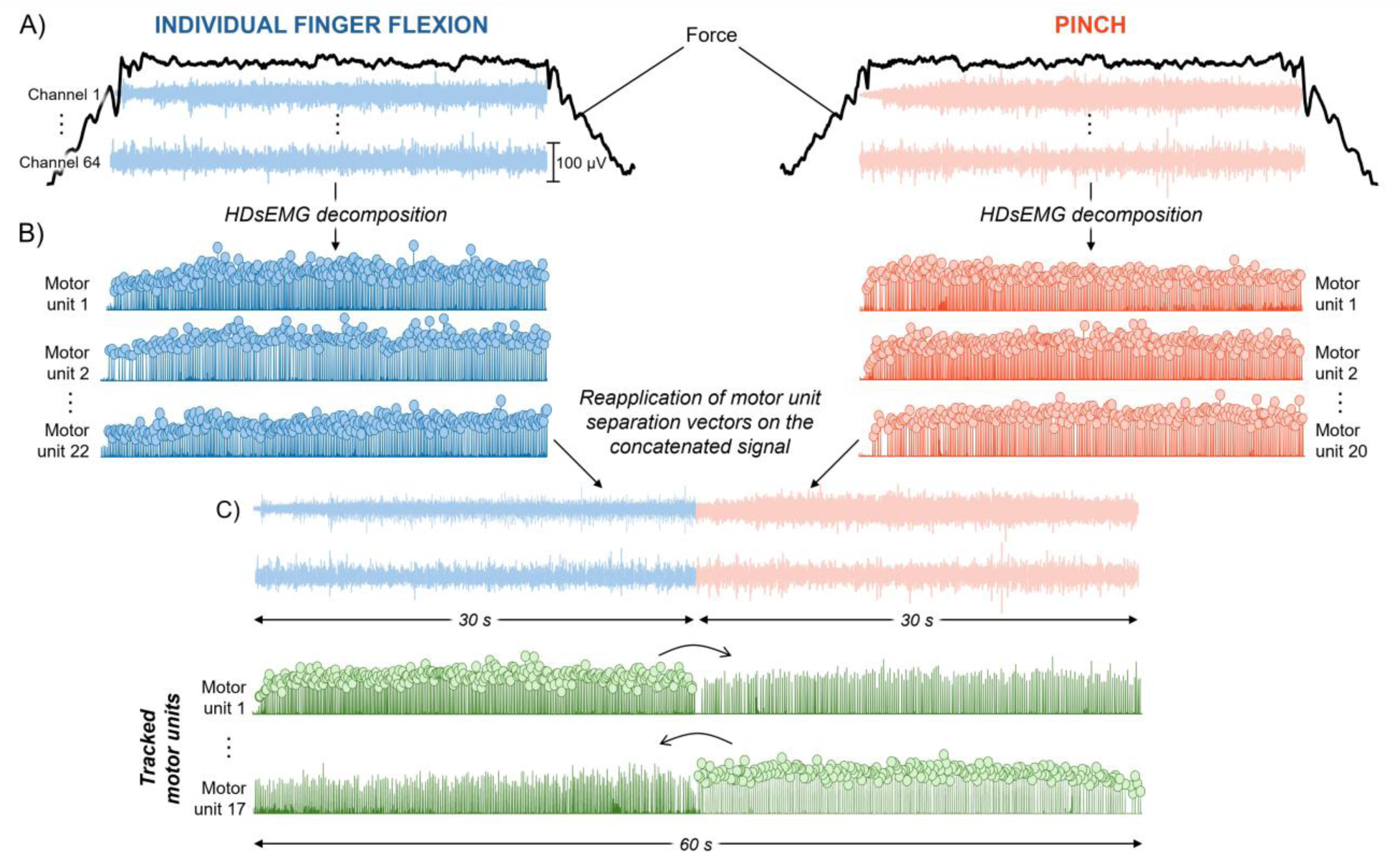
A) High-density surface EMG (HDsEMG) signals recorded during individual index finger flexion (blue, left), and pinch (orange, right) were decomposed into individual motor unit spike trains using a convolutive blind source separation approach. B) To ensure that identical motor units were examined across conditions, the separation vectors identified in each task were reapplied to the concatenated HDsEMGs of both tasks. This procedure allowed consistent tracking of motor units between individual index finger flexion and pinch tasks. Representative examples of the resulting innervation pulse trains are illustrated. C) Tracked motor units after reapplication of separation vector on the concatenated HDsEMGs are shown in green.

#### Coherence analysis

To estimate the shared synaptic inputs to motor unit spike trains, we used coherence analysis (Negro & Farina, 2012; Dideriksen et al., 2018). Coherence is a frequency-domain measure of the linear correlation between two signals. In the context of motor unit analysis, it reflects, for each frequency, the degree of common oscillations in the motoneuron pool and it is often used to infer shared synaptic inputs during voluntary contractions (Halliday et al., 1995, Castronovo et al., 2015). For this analysis, motor unit discharge times were converted into binary spike trains, with 1 indicating the presence of a discharge time and 0 its absence. To increase reliability and reduce the influence of single motor units, we estimated coherence between cumulative spike trains (CSTs) formed from random, non-overlapping subsets of the identified units. Specifically, for participants with at least 4 matched motor units, two disjoint groups of motor units were randomly selected (each group containing 50% of the identified units), their spike trains were summed to create two CSTs, and coherence between these CSTs was computed using a cross-power spectral density approach. This procedure was repeated for 100 iterations and the average coherence across iterations (i.e., pooled coherence) was considered. For all iterations, coherence was calculated between the two detrended CSTs using a 1-s Hanning window with approximately 0.95 s overlap and a spectral resolution of ten times the sampling frequency. The pooled coherence was then transformed into standardized z-scores following the approach outlined by Gallet and Julien (2011). Only z-coherence values exceeding the bias, defined as the average z-coherence within the 250–500 Hz range (Castronovo et al., 2015), were included for further analysis (Figure 3B). The area under the curve of z-coherence was calculated in the delta (1-5 Hz), alpha (5-15 Hz), and beta (15-35 Hz) bands, which are associated with different physiological information. The delta band is generally attributed to descending voluntary commands and reflects the common drive to motoneurons during steady contractions (Negro et al., 2015). The alpha band may reflect activity within spinal or subcortical circuits, including reflex pathways and proprioceptive feedback mechanisms (Boonstra et al., 2012). In contrast, the beta band is associated with cortical contributions, particularly corticospinal drive (McManus et al., 2019).

**Figure 3.**
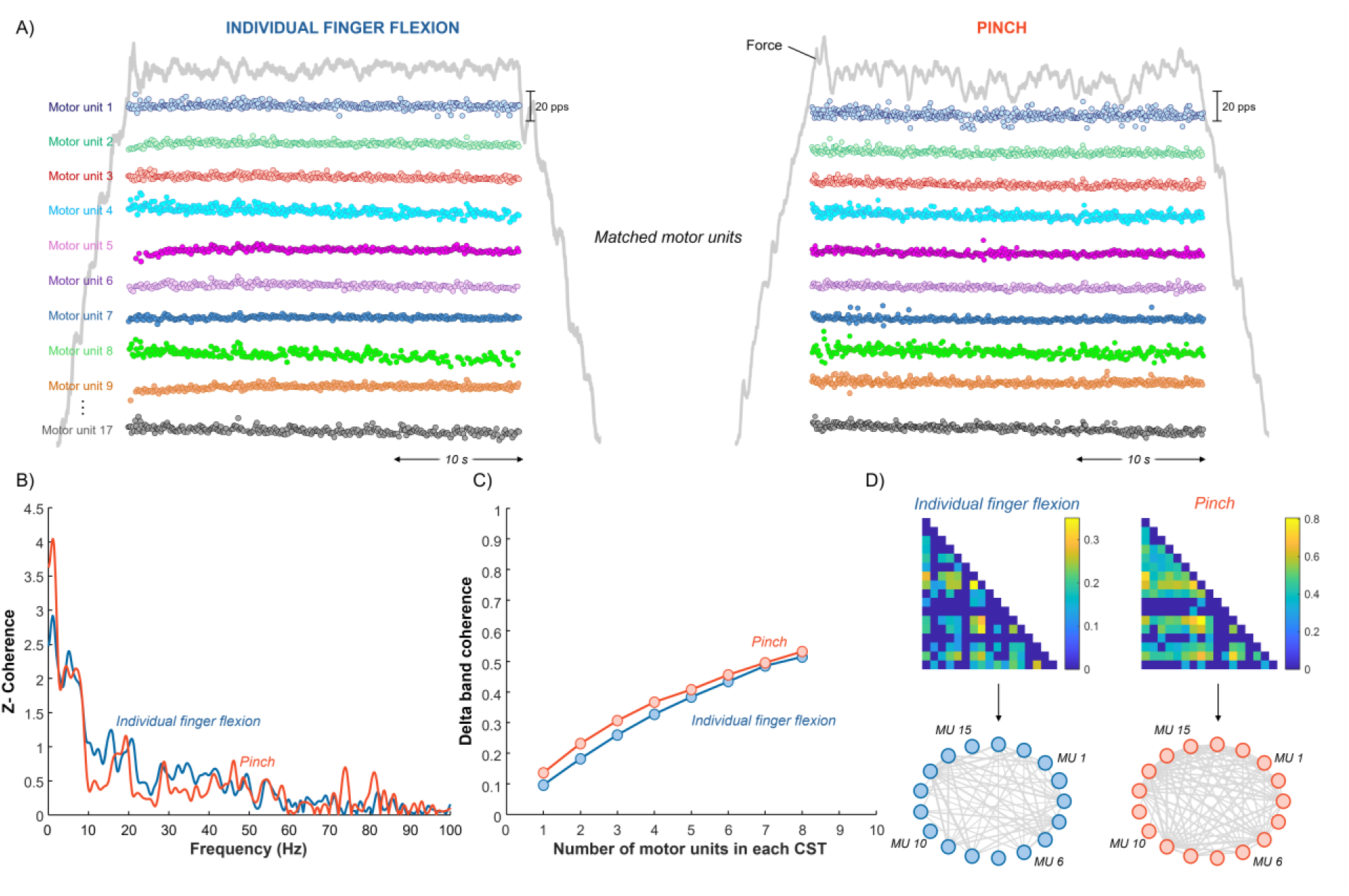
Representative contractions recorded at 20% MVC in both individual index finger flexion and pinch tasks. A) instantaneous discharge rates of matched motor units across tasks and concurrent force trace. Each motor unit is shown in a different color. B) z-coherence curves calculated from spectrum motor units shown in A). Individual index finger flexion and pinch tasks are represented by blue and orange traces, respectively. C) Curves of mean delta-band pooled coherence as a function of the number of motor units included in the cumulative spike train (CST), which were used to estimate the proportion of common input (PCI) index. Individual index finger flexion and pinch tasks are represented by blue and orange traces, respectively. D) Mutual Information showing the number of nodes and the way they are connected to each other in both individual index finger flexion and pinch tasks.

#### Proportion of common input (PCI) index

To quantify the extent of common synaptic input with respect to independent one across the motoneuron pool, we computed the Proportion of Common Input (PCI) index, as described by Negro et al. (2016). The PCI index estimates the fraction of total input variance that can be attributed to common synaptic drive across a pool of motoneurons. For this, motor unit spike trains were converted to binary discharge trains and used to build CSTs. For each condition, CSTs were formed from two equal, non–overlapping subsets of units. The subset size is expressed as *m* (motor units per CST), with *m* ranging from 1 to *N/2*, where *N* is the number of matched motor units. For each *m*, up to 100 random permutations were performed. The mean coherence in the delta band (1–5 Hz) was taken as the coherence value for that permutation, and values were averaged across all permutations to produce the delta band coherence vs. *m* curve (Figure 3C). The resulting curve was fitted with a nonlinear model (see equation 4 in Negro et al. 2016). Then, the model parameters A and B were estimated by nonlinear least squares, and the PCI was computed as *PCI = sqrt(A/B)*. Higher PCI values indicate a greater proportion of shared input among motor units, reflecting stronger common drive (Figure 3C).

#### Network-information framework

To characterize non-linear interactions across motor units, we applied the network-information theoretic framework of O’Reilly and Delis (2022, 2024). This framework has been recently applied to investigate the non-linear associations between motor units (Cabral, et al., 2025). First, smoothed discharge rates were obtained by convolving binary spike trains with a 400-ms Hanning window and high-pass filtering at 0.75 Hz using a third-order Butterworth filter to eliminate trends (De Luca et al., 1982). Pairwise mutual information (MI) was then estimated for every unique motor unit pair. This produced a symmetric adjacency matrix that captured the functional connectivity across motor units (Figure 3D). To isolate meaningful connections, we applied a modified percolation analysis (Gallos et al., 2012), which determined a threshold, the percolation point, beyond which associations were considered significant. These connections were then mapped into a motor unit network using graph-theoretical constructs, where each motor unit was represented as a node and each significant link as an edge (Figure 3D). For visualization, we adopted a circular layout to display the matched motor units across tasks. From this network we computed the weighted density, defined as the sum of all edge weights in the network divided by the maximum possible number of edges. This metric reflects how strong the network links are overall (i.e., strength of the network) (Ioannis et al., 2007).

### Statistical analysis

The statistical approach utilized a linear mixed model (LMM) that was applied using R (version 4.x) with the *lme4* and *lmerTest* packages, implemented within the Jamovi statistical software (version 2.3.28.0). LMMs were used to compare the z-coherence in the delta, alpha and beta bands, and the PCI values between individual index finger flexion and pinch tasks. LMMs were also used to compare the weighted density and proportion of motor units in the first low-dimensional component. For all outcomes, random intercepts models were used with task (individual index finger flexion and pinch) and force level (10% and 20% MVC) as fixed effects, and participant as a random effect, with additional random intercepts for the subject-by-task and subject-by-force interactions. Degrees of freedom and p-values for fixed effects were estimated using Satterthwaite’s method, and estimated marginal means were computed using the Kenward-Roger method. In cases of significant interactions, Holm’s post hoc tests were performed for pairwise comparisons. Statistical significance was set as α = 0.05. Results are reported as mean ± standard deviation in the text and tables. All individual data on matched motor unit discharge times are available at https://doi.org/10.6084/m9.figshare.31971636.

## RESULTS

### Motor unit yield

To investigate the changes in neural control of FDI motor units between individual index finger flexion and pinch tasks, HDsEMG were decomposed into motor unit spike trains and tracked across tasks. **Table 1** contains the average number of motor units identified and matched per participant. The average SIL value for matched units across tasks was 0.92 ± 0.02 for 10% and 0.91 ± 0.01 for 20% MVC.

**Table 1.** The average number of motor units per participant, separately per task.

|  | 10% MVC | 20% MVC |
| --- | --- | --- |
| **Individual index finger flexion** | $18 \pm 4$ | $20 \pm 4$ |
| **Pinch** | $19 \pm 5$ | $19 \pm 6$ |
| **Matched across tasks** | $10 \pm 4$ | $13 \pm 3$ |

### Coherence analysis

To investigate changes in shared synaptic inputs to FDI motor units between individual index finger flexion and pinch tasks, coherence analysis between motor unit spike trains was performed with focus on the delta, alpha and beta bands. This analysis was carried out in 13 out of 16 subjects, as one subject did not have at least four matched motor units (see Methods) and two outliers were removed based on the interquartile range method. Figure 4A is a representative example at 20% MVC, with substantial overlap between the coherence spectra of individual index finger flexion and pinch tasks. No clear differences were observed across the delta, alpha, and beta frequency bands. In agreement with the representative case, the group results revealed no statistical differences between tasks for either a task main effect or a task × force interaction for the area under the curve of the delta band (Figure 4C; LMM; Task main effect: F(1, 16.06) = 2.15, P = 0.161; Task × Force interaction: F(1, 14.27) = 0.05, P = 0.823), alpha band (Figure 4D; LMM; Task main effect: F(1, 24.00) = 3.31, P = 0.081; Task × Force interaction: F(1, 24.00) = 0.32, P = 0.576), or beta band coherence (Figure 4E; LMM; Task main effect: F(1, 13.72) = 0.11, P = 0.745; Task × Force interaction: F(1, 9.59) = 4.33, P = 0.065).

**Figure 4.**
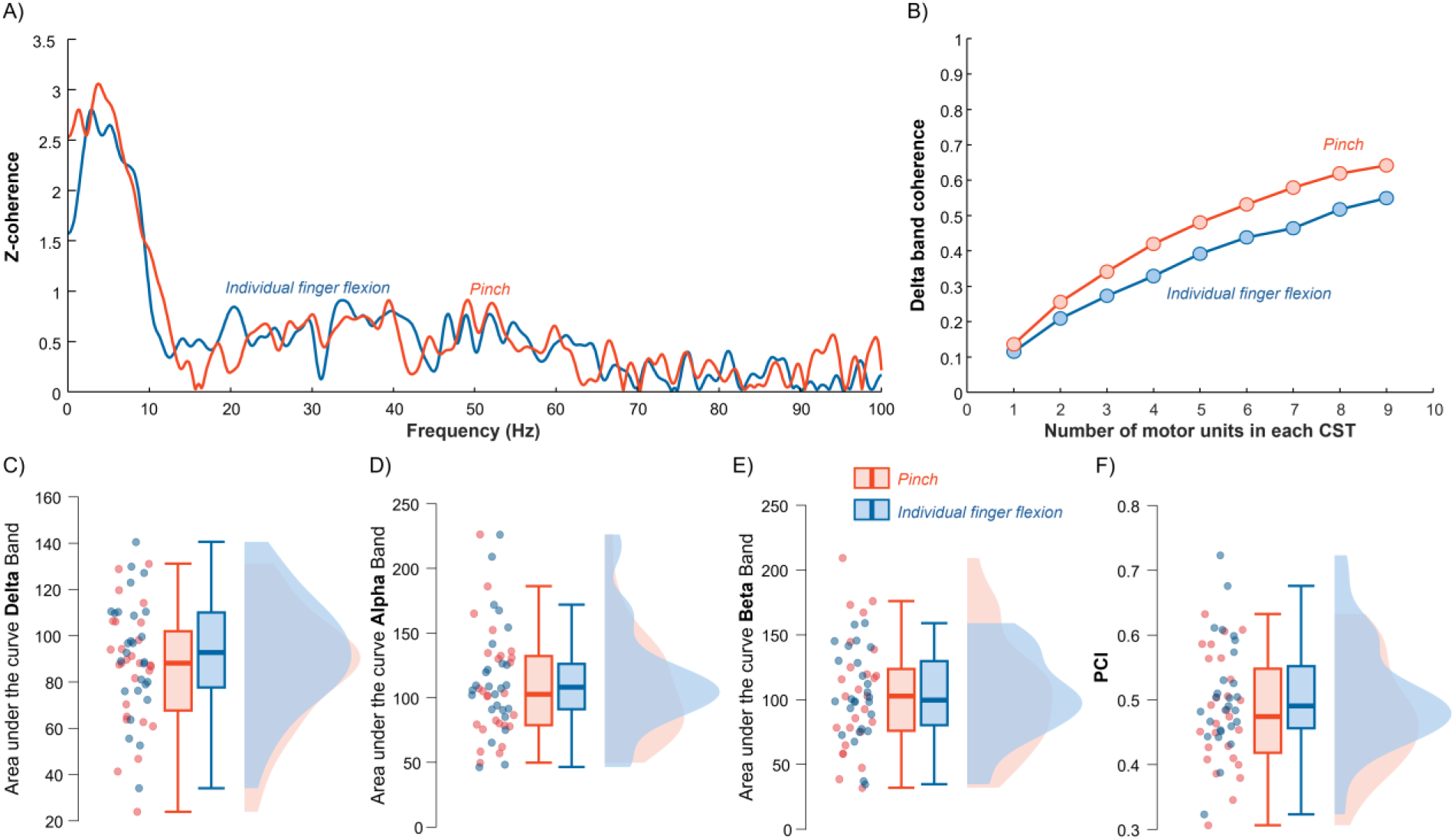
Coherence (A) and PCI (B) analysis from FDI motor units for a representative participant. Motor units were tracked between individual index finger flexion (blue) and pinch (orange) tasks. Group results for delta band z-coherence (C), alpha band z-coherence (D), beta band z-coherence (E), and Proportion of Common Input (PCI) index (F). Circles represent individual participants. Data are represented as median (horizontal lines), interquartile range (boxes), and distribution range (whiskers).

### Proportion of common input (PCI) index

The PCI index was computed to quantify the relative contribution of common versus independent synaptic inputs across the motoneuron pool. After computing the average delta-band coherence as a function of the number of motor units included in the CST, the PCI index was calculated as an estimate of the overall level of common input reflected in the coherence–motor unit number relationship. Figure 4B illustrates the relationship between average delta-band coherence and the number of motor units for the same participant shown in Figure 4A. Coherence increased as additional motor units were incorporated into the CST; however, the rate of increase was nearly identical for the individual index finger flexion and pinch tasks. Similarly to the representative case, there were no statistical differences in PCI index between tasks and across forces (Figure 4F; LMM; Task main effect: F(1, 16.0) = 1.50, P = 0.240; Task × Force: F(1, 13.8) = 0.05, P = 0.820).

### Network-information framework

To assess the non-linear relationship between pairs of motor unit smoothed discharge rates, we characterized motor unit networks across tasks using a network-information framework. Figure 5A presents a representative example at 20% MVC, illustrating the motor unit network connectivity. Compared to the individual index finger flexion task, the pinch condition exhibits a visibly denser network, characterized by a greater number of connections between nodes and stronger overall coupling. Indeed, the group results revealed a significant increase in weighted density during pinch compared to individual index finger flexion (Figure 5B; LMM; Task main effect: F(1, 17.75) = 7.55, P = 0.013), independent of the force level (Task × Force: F(1, 9.94) = 0.37, P = 0.556).

**Figure 5.**
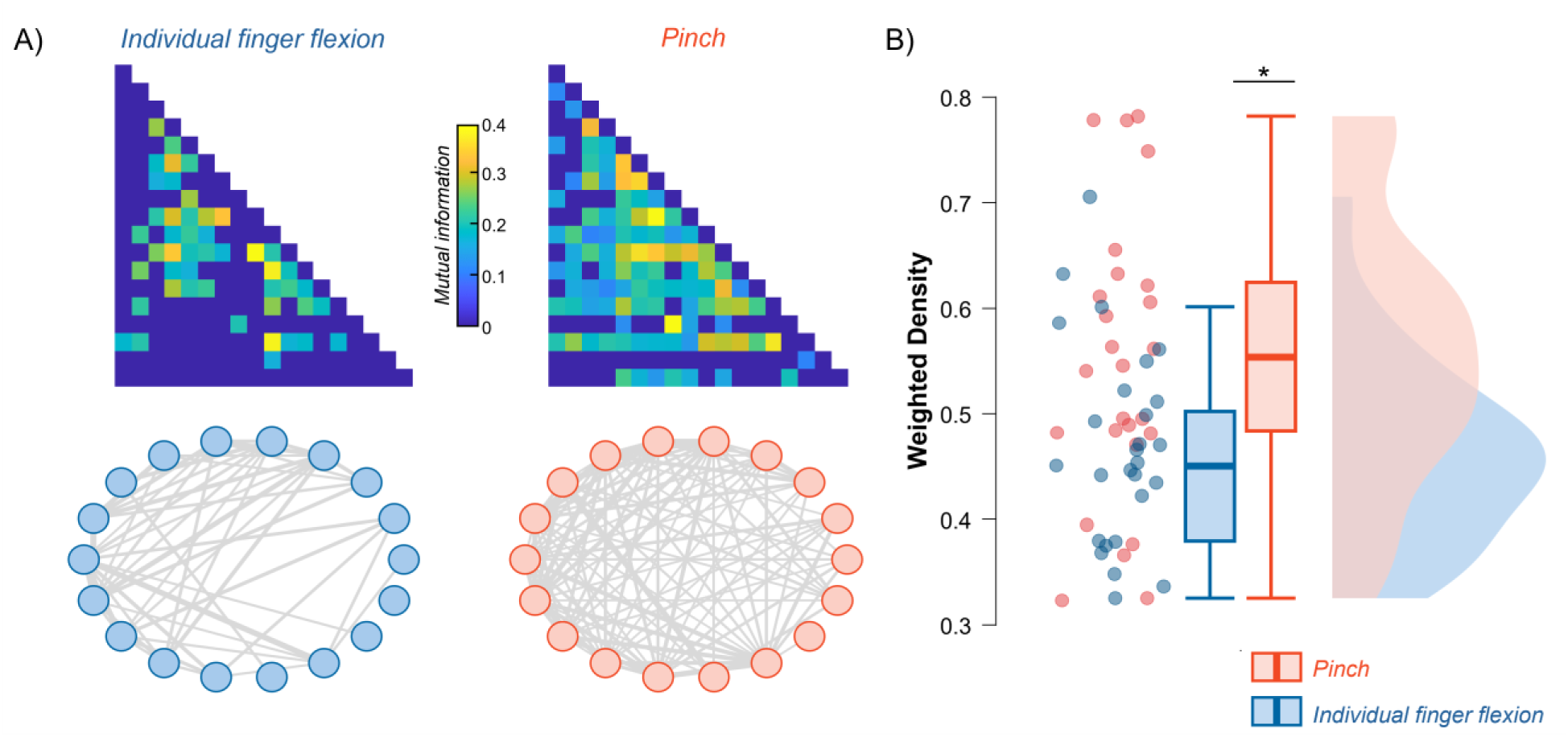
A) Thresholded mutual-information (MI) adjacency matrices after Gaussian-copula normalization (top panel). Only values above the modified percolation threshold are displayed (color bar indicates MI magnitude). Corresponding graph representations of motor unit coupling using a circular layout are shown in the bottom panel. Each node represents a matched motor unit and each edge a significant MI link whose thickness is proportional to weight. Identical node order and edge-weight scaling were used to allow direct visual comparison of connection strength and topology. B) Group results for the weight density of motoneuron networks for individual index finger flexion (orange) and pinch (blue) tasks. Data are represented as median (horizontal lines), interquartile range (boxes), and distribution range (whiskers).

## DISCUSSION

This study investigated the neural coupling between motor unit spike trains of the FDI muscle during isolated index finger flexion and precision pinch tasks performed at two matched force levels (10% and 20% MVC). By combining second-order correlation methods (coherence analysis and Proportion of Common Input index) with a network-information framework based on mutual information, we assessed whether common synaptic inputs to the same motoneuron pool adapt as a function of task demands. The main findings reveal a dissociation between linear and nonlinear measures of motor unit coupling. Coherence-based analysis showed no significant differences between tasks, whereas network density derived from mutual information revealed significantly stronger nonlinear coupling during the pinch task than during isolated index finger flexion. These results suggest that precision pinch tasks rely on stronger higher-order common inputs and distinct neural control strategies that are not fully captured by traditional linear measures. These findings emphasize the importance of nonlinear interactions in coordinating motor unit activity during functionally relevant fine motor control.

The coherence analysis and PCI index resulted in no significant differences in common synaptic input between isolated index finger flexion and pinch tasks across the delta, alpha, and beta frequency bands. This finding was unexpected given the substantial differences in task complexity and coordination demands, particularly in light of recent studies demonstrating task-dependent modulation of common inputs in hand muscles using similar linear methods (Tanzarella et al. 2021, Cabral et al. 2025, Del Vecchio et al. 2022). In extrinsic hand muscles, current literature suggests that task demands actively modulate shared synaptic inputs. Tanzarella et al. (2021) reported that motoneuron synergies differ systematically across grip types, with low-dimensional components reorganizing depending on involved fingers. Similarly, Cabral et al. (2025) stated that alpha-band coherence across extrinsic hand muscles decreases during grasping compared to four-finger flexion, while Del Vecchio et al. (2022) suggested that common inputs vary across motor neurons in extrinsic muscles in a task-dependent manner.

In contrast, our results indicate that the FDI, an intrinsic hand muscle, maintains stable linear common input across tasks when force levels are matched. The delta band (1-5 Hz) mirrors common drive to motoneurons during steady contractions (Negro et al., 2016; Conway et al., 1995), while the beta band (15-35 Hz) reflects corticospinal drive (McManus et al., 2019; Baker et al., 1997). The absence of task-dependent modulation suggests that the descending cortical drive, as reflected in linear oscillatory coupling, remains relatively stable, with the FDI generating similar force levels whether acting in isolation or synergistically with the thenar during pinch. The dichotomy between extrinsic and intrinsic hand muscles reflects fundamental differences in their neural control architecture. Intrinsic muscles, like the FDI, exhibit greater functional independence and receive more selective and independent descending inputs compared to extrinsic muscles (McIsaac & Fuglevand, 2008; Winges et al., 2008). McIsaac and Fuglevand (2008) demonstrated that motor unit synchrony across intrinsic muscles during precision grip is markedly lower than in extrinsic muscles, with Common Input Strength analysis indicating muscle-specific rather than shared synaptic inputs. This functional independence may explain why the FDI maintains stable baseline corticospinal drive across tasks, while extrinsic muscles exhibit pronounced task-dependent reorganization. Moreover, corticospinal beta oscillations are transmitted uniformly across the motoneuron pool regardless of recruitment threshold (Abbagnano et al., 2025), suggesting that the transmission pathway itself maintains stable properties across different motor contexts. This may further contribute to the absence of task-dependent differences in beta coherence. Finally, as coherence captures only linear relations, it may fail to capture the nonlinear influences of beta-band inputs on force control arising from the motoneuron pool’s inherent nonlinearities (Negro et al., 2016), underscoring the value of complementary analyses, such as Mutual Information, to fully characterize motor unit coupling.

In contrast to the above discussed linear analysis techniques, the network-information framework revealed significantly increased weighted density during the pinch task compared to isolated index finger flexion. This increase in network density indicates stronger, more numerous nonlinear associations between motor unit discharge patterns when the FDI acts synergistically with the thenar during precision grip, compared with index finger flexion alone. Schieber’s concept of “individuated” movement (1990) provides a useful perspective: independent finger movements may require suppression of the hand’s intrinsic tendency to act as a coordinated unit. In this context, pinch represents a functionally integrated synergy rather than an isolated action, potentially allowing higher-order interactions among motor-unit discharge patterns and their shared synaptic inputs to emerge more strongly. Mutual information captures these statistical dependencies beyond linear correlations that may arise from complex synaptic integration, proprioceptive feedback-driven modulation, and intrinsic motoneuron nonlinearities (O’Reilly & Delis, 2022, 2024). The dissociation observed in this study, with stable linear coherence alongside increased nonlinear network density, suggests that precision grip engages neural coordination mechanisms that operate through nonlinear pathways not identified by conventional coherence measures.

Several neurophysiological factors may contribute to this enhanced nonlinear coupling. First, precision pinch involves stronger higher-order common inputs compared with isolated FDI flexion. This possibility aligns with previous evidence indicating increased corticospinal excitability and enhanced motor cortical engagement during precision grip compared to simple finger flexion (Tinazzi et al., 2003). Second, proprioceptive feedback may play a role in the increased non-linear interaction between spinal motoneurons. During pinch, mechanical interaction between the thumb and index finger generates complex proprioceptive and tactile inputs from muscle spindles, Golgi tendon organs, and cutaneous receptors (Bourguignon et al., 2015). Such multisensory integration may introduce nonlinear dependencies between motor unit discharge patterns. While proprioceptive feedback can decorrelate motor unit discharge rates through discordant excitation of homonymous motoneurons (De Luca & Kline, 2011), it may simultaneously increase higher-order or nonlinear coupling within the motor unit network. Third, the pinch task requires coordinated activation of the FDI and thenar muscles, which are innervated by different nerves but must generate opposing forces to stabilize thumb-index contact. Even though thenar activity was not recorded in the present study, its involvement may increase the complexity of common synaptic inputs received by the FDI motoneuron pool. This interpretation is supported by evidence of heteronymous H-reflex pathways between median-innervated thenar muscles and FDI motor neurons, indicating shared afferent modulation across these pools (Baudry et al., 2009).

### Functional and translational implications

The increased nonlinear coupling during pinch has important functional implications for understanding precision grip control. Precision grip requires not only adequate force production but also fine temporal coordination, stability against perturbations, and rapid adaptation to sensory feedback (Johansson & Flanagan, 2009). Nonlinear mechanisms may facilitate integration across multiple temporal scales, allowing fast cortical oscillations (e.g., beta band activity) to influence slower force modulation through nonlinear transformations within the motoneuron pool (Watanabe & Kohn 2015). Recent studies using cross-frequency coherence and n:m coherence have demonstrated that nonlinear corticomuscular coupling exists during various motor tasks, with different frequency bands interacting in complex ways that are not captured by traditional linear coherence (Yang et al., 2016). Our results advance these findings by demonstrating that nonlinear coupling at the motor unit level is modulated by task demands, suggesting that the nervous system actively regulates the balance between linear and nonlinear control mechanisms depending on functional requirements. This perspective aligns with Schieber’s (1990) population-coding framework, in which cortical control of movement emerges from distributed neuronal populations rather than independent commands to individual muscles. Therefore, the output reaching the motoneuron pool represents a compressed summary of complex cortical processing where linear coherence may reflect the shared rhythm component of this output, and the nonlinear network density may capture higher-order structure within the population code that supports synergy control.

Understanding linear and nonlinear motor unit coupling has important implications for rehabilitation after neurological injury. Impaired pinch control following stroke may arise not only from reduced corticospinal drive but also from disrupted nonlinear integration involving proprioceptive and spinal circuits. Brain-machine interface (BMI) systems that reinforce sensorimotor contingency and enhance proprioceptive feedback can reorganize motor assemblies (Caria et al., 2020) and could potentially incorporate network density metrics as feedback signals to target higher-order coordination patterns during precision grip training. Moreover, the dissociation between linear coherence and nonlinear network density may provide complementary biomarkers of recovery.

## LIMITATIONS AND CONCLUSIONS

There are a few limitations in this study that need to be acknowledged. First, we analyzed motor units exclusively from the FDI muscle and did not analyze the thenar, which limits direct inference about intermuscular coupling during pinch. Future studies incorporating motor units from both FDI and thenar muscles could construct integrated networks to explicitly quantify intra- and intermuscular coupling during precision grip. Second, we examined only two submaximal force levels (10% and 20% MVC). The balance between linear and nonlinear coupling may differ at higher force levels, where most motor units are already recruited and force adjustments rely more heavily on rate coding. Extending the analysis across a wider force range would clarify whether the dissociation between coherence and nonlinear network density persists at higher contraction levels. Finally, it is important to note that coherence analysis is inherently limited to detecting linear relations between signals. The absence of task-dependent differences in coherence does not necessarily imply a lack of modulation in descending cortical drive, but rather it indicates that any such modulation does not manifest as increased linear oscillatory synchronization within the analyzed frequency bands. To best address these limitations, which is the key strength of this study, the integration of complementary analytical approaches to characterize motor unit coupling was implemented. By combining coherence analysis (capturing linear oscillatory coupling), PCI index (estimating the balance between shared and independent inputs), and mutual information-based network analysis (capturing nonlinear statistical dependencies), we provide a more comprehensive characterization of motor unit coordination than would be possible with any single method alone. This multimethod framework directly addresses limitations of earlier studies that relied exclusively on pairwise linear correlation methods, which can be biased by discharge statistics and may underestimate complex dependencies (de la Rocha et al., 2007; Farina & Negro 2015).

In conclusion, this study demonstrates that precision pinch engages distinct neural control strategies, characterized by preserved linear common input yet markedly enhanced nonlinear coupling among FDI motor units. By reporting stronger higher-order interactions during pinch than during isolated index-finger flexion, these findings demonstrate that precision grip involves stronger shared inputs distributed across the entire motoneuron pool than individual finger contractions. Interestingly, these enhanced interactions operate through neural processes that extend beyond pairwise associations and are not fully captured by traditional linear methods. The dissociation between linear coherence measures (which remained stable across tasks) and nonlinear network density (which increased significantly during pinch) indicates that functionally relevant precision grip tasks recruit additional tiers of neural coordination that rely on complex, nonlinear mechanisms. This dissociation between linear and nonlinear approaches accentuates the importance of combining complementary analytical tools, integrating both second-order correlation methods and mutual information-based network analysis, to fully characterize motor unit control. Future work incorporating simultaneous recordings from multiple hand muscles across broader force ranges and task contexts will help clarify how the nervous system balances linear and nonlinear control mechanisms to achieve the remarkable dimensionality and precision of human hand function.

## ACKNOWLEDGEMENTS

This study was funded by the European Research Council Consolidator Grant INcEPTION contract 21 no. 101045605. J Greig Inglis was supported by the Marie Skłodowska-Curie Actions Grant 22 ‘MUDecomp’ agreement no. 101151712.

## References

1. Abbagnano, E., Pascual-Valdunciel, A., Zicher, B., Ibañez, J., & Farina, D. (2025). Projection of cortical beta band oscillations to a motor neuron pool across the full range of recruitment. Journal of Neuroscience, 45(31).

2. Ahn, Y. Y., Bagrow, J. P., & Lehmann, S. (2010). Link communities reveal multiscale complexity in networks. nature, 466(7307), 761–764.

3. Amjad, A. M., Halliday, D. M., Rosenberg, J. R., & Conway, B. A. (1997). An extended difference of coherence test for comparing and combining several independent coherence estimates: Theory and application to the study of motor units and physiological tremor. Journal of Neuroscience Methods, 73(1), 69–79.

4. Baker, S. N., Olivier, E., & Lemon, R. N. (1997). Coherent oscillations in monkey motor cortex and hand muscle EMG show task-dependent modulation. The Journal of physiology, 501(Pt 1), 225.

5. Bizzi, E., & Cheung, V. C. (2013). The neural origin of muscle synergies. Frontiers in computational neuroscience, 7, 51.

6. Boonstra, T. W., & Breakspear, M. (2012). Neural mechanisms of intermuscular coherence: implications for the rectification of surface electromyography. Journal of neurophysiology, 107(3), 796–807.

7. Baudry, S., Jordan, K., & Enoka, R. M. (2009). Heteronymous reflex responses in a hand muscle when maintaining constant finger force or position at different contraction intensities. Clinical neurophysiology, 120(1), 210–217.

8. Bourguignon, M., Piitulainen, H., De Tiège, X., Jousmäki, V., & Hari, R. (2015). Corticokinematic coherence mainly reflects movement-induced proprioceptive feedback. Neuroimage, 106, 382–390.

9. Cabral, H. V., Cudicio, A., Bonardi, A., Del Vecchio, A., Falciati, L., Orizio, C., Martinez-Valdes, E., & Negro, F. (2024). Neural filtering of physiological tremor oscillations to spinal motor neurons mediates short-term acquisition of a skill learning task. eNeuro, 11(7). 10.1523/ENEURO.0043-24.2024

10. Cabral, H. V., Inglis, J. G., Pourreza, E., Dos Santos, M. A., Cosentino, C., O’Reilly, D., … & Negro, F. (2025). A single low-dimensional neural component of spinal motor neuron activity explains force generation across repetitive isometric tasks. iScience, 28(10).

11. Cabral, H. V., Cosentino, C., Rizzardi, A., Inglis, J. G., Fuglevand, A. J., & Negro, F. (2025). Adaptations in common synaptic inputs to spinal motor neurons during grasping versus a less functional hand task. Journal of Applied Physiology, 139(3), 776–786.

12. Caria, A., da Rocha, J. L. D., Gallitto, G., Birbaumer, N., Sitaram, R., & Murguialday, A. R. (2020). Brain–machine interface induced morpho-functional remodeling of the neural motor system in severe chronic stroke. Neurotherapeutics, 17(2), 635–650.

13. Castronovo, A. M., Negro, F., Conforto, S., & Farina, D. (2015). The proportion of common synaptic input to motor neurons increases with an increase in net excitatory input. Journal of Applied Physiology, 119(11). 10.1152/japplphysiol.00255.2015

14. Chen, M., & Zhou, P. (2022). Caution is necessary for acceptance of motor units with intermediate matching in surface EMG decomposition. Frontiers in Neuroscience, 16, 876659.

15. Conway, B. A., Halliday, D. M., Farmer, S. F., Shahani, U., Maas, P., Weir, A. I., & Rosenberg, J. R. (1995). Synchronization between motor cortex and spinal motoneuronal pool during the performance of a maintained motor task in man. The Journal of physiology, 489(3), 917–924.

16. Cooney III, W. P., An, K. N., Daube, J. R., & Askew, L. J. (1985). Electromyographic analysis of the thumb: a study of isometric forces in pinch and grasp. The Journal of hand surgery, 10(2), 202–210.

17. Dai, C., & Hu, X. (2019). Independent component analysis-based algorithms for high-density electromyogram decomposition: Systematic evaluation through simulation. Computers in Biology and Medicine, 109, 171–181.

18. De La Rocha, J., Doiron, B., Shea-Brown, E., Josić, K., & Reyes, A. (2007). Correlation between neural spike trains increases with firing rate. Nature, 448(7155), 802–806.

19. De Luca, C. J., LeFever, R. S., McCue, M. P., & Xenakis, A. P. (1982). Control scheme governing concurrently active human motor units during voluntary contractions. The Journal of physiology, 329(1), 129–142.

20. De Luca, C. J., & Kline, J. C. (2011). Influence of proprioceptive feedback on the firing rate and recruitment of motoneurons. Journal of Neural Engineering, 9(1), 016007.

21. Del Vecchio, A., Germer, C. M., Elias, L. A., Fu, Q., Fine, J., Santello, M., & Farina, D. (2019). The human central nervous system transmits common synaptic inputs to distinct motor neuron pools during non-synergistic digit actions. The Journal of Physiology, 597(24), 5935–5948.

22. Del Vecchio, A., Germer, C. M., Kinfe, T. M., Nuccio, S., Hug, F., Eskofier, B., … & Enoka, R. M. (2023). The forces generated by agonist muscles during isometric contractions arise from motor unit synergies. Journal of Neuroscience, 43(16), 2860–2873.

23. de Vries, I. E., Daffertshofer, A., Stegeman, D. F., & Boonstra, T. W. (2016). Functional connectivity in the neuromuscular system underlying bimanual coordination. Journal of neurophysiology, 116(6), 2576–2585.

24. Dideriksen, J. L., Negro, F., Falla, D., Kristensen, S. R., Mrachacz-Kersting, N., & Farina, D. (2018). Coherence of the surface EMG and common synaptic input to motor neurons. Frontiers in Human Neuroscience, 12, 207.

25. Farina, D., Negro, F., & Dideriksen, J. L. (2014). The effective neural drive to muscles is the common synaptic input to motor neurons. The Journal of physiology, 592(16), 3427–3441.

26. Farina, D., & Negro, F. (2015). Common synaptic input to motor neurons, motor unit synchronization, and force control. Exercise and sport sciences reviews, 43(1), 23–33.

27. Fuglevand, A. J. (2011). Mechanical properties and neural control of human hand motor units. The Journal of physiology, 589(23), 5595–5602.

28. Gallet C, Julien C (2011) The significance threshold for coherence when using the Welch’s periodogram method: effect of overlapping segments. Biomed Signal Process Control 6:405–409.

29. Gallos, L.K., Makse, H.A., and Sigman, M. (2012). A small world of weak ties provides 897 optimal global integration of self-similar modules in functional brain networks. 898 Proceedings of the National Academy of Sciences 109, 2825–2830. 899 doi:10.1073/pnas.1106612109

30. Gentner, R., & Classen, J. (2006). Modular organization of finger movements by the human central nervous system. Neuron, 52(4), 731–742.

31. Giurintano, D. J., Hollister, A. M., Buford, W. L., Thompson, D. E., & Myers, L. M. (1995). A virtual five-link model of the thumb. Medical engineering & physics, 17(4), 297–303.

32. Halliday, D. M., Rosenberg, J. R., Amjad, A. M., Breeze, P., Conway, B. A., & Farmer, S. F. (1995). A framework for the analysis of mixed time series/point process data theory and application to the study of physiological tremor, single motor unit discharges and electromyograms. Progress in biophysics and molecular biology, 64(2-3), 237–278.

33. Hug, F., Avrillon, S., Del Vecchio, A., Casolo, A., Ibanez, J., Nuccio, S., Rossato, J., Holobar, A., & Farina, D. (2021). Analysis of motor unit spike trains estimated from high-density surface electromyography is highly reliable across operators. Journal of Electromyography and Kinesiology, 58, 102548.

34. Hug, F., Avrillon, S., Ibáñez, J., & Farina, D. (2023). Common synaptic input, synergies and size principle: Control of spinal motor neurons for movement generation. The Journal of physiology, 601(1), 11–20.

35. Hu, N., Cabral, H. V., Pourreza, E., Inglis, J. G., Desmons, M., & Negro, F. (2025). Motor unit rate coding in intrinsic hand muscles during isolated finger contractions and pinch task. bioRxiv, 2025-12.

36. Ioannis, A., & Eleni, T. (2007). Statistical analysis of weighted networks. arXiv preprint arXiv:0704.0686.

37. Johansson, R. S., Flanagan, J. R., & Johansson, R. S. (2009). Sensory control of object manipulation. Sensorimotor control of grasping: Physiology and pathophysiology, 141–160.

38. Kozin, S. H., Porter, S., Clark, P., & Thoder, J. J. (1999). The contribution of the intrinsic muscles to grip and pinch strength. The Journal of hand surgery, 24(1), 64–72.

39. Laine, C. M., & Valero-Cuevas, F. J. (2017). Intermuscular coherence reflects functional coordination. Journal of Neurophysiology, 118(3), 1775–1787. 10.1152/jn.00204.2017

40. Latash, M. L. (2012). The bliss (not the problem) of motor abundance (not redundancy). Experimental brain research, 217(1), 1–5.

41. Lawrence, D. G., & Kuypers, H. G. (1968). The functional organization of the motor system in the monkey: I. The effects of bilateral pyramidal lesions. Brain, 91(1), 1–14.

42. Lawrence, D. G., & Hopkins, D. A. (1976). The development of motor control in the rhesus monkey: evidence concerning the role of corticomotoneuronal connections. Brain: a journal of neurology, 99(2), 235–254.

43. Lemon, R. N., Mantel, G. W., & Muir, R. B. (1986). Corticospinal facilitation of hand muscles during voluntary movement in the conscious monkey. The Journal of physiology, 381(1), 497–527.

44. Lemon, R. N., Johansson, R. S., & Westling, G. (1995). Corticospinal control during reach, grasp, and precision lift in man. Journal of Neuroscience, 15(9), 6145–6156.

45. Liu, Y., & Rouiller, E. M. (1999). Mechanisms of recovery of dexterity following unilateral lesion of the sensorimotor cortex in adult monkeys. Experimental brain research, 128(1), 149–159.

46. Martinez-Valdes, E., & Negro, F. (2023). Neuromuscular function: High-density surface electromyography. In Neuromuscular Assessments of Form and Function (pp. 105–123). New York, NY: Springer US.

47. McManus, L., Flood, M. W., & Lowery, M. M. (2019). Beta-band motor unit coherence and nonlinear surface EMG features of the first dorsal interosseous muscle vary with force. Journal of Neurophysiology, 122(3), 1147–1162.

48. Maillet, J., Avrillon, S., Nordez, A., Rossi, J., & Hug, F. (2022). Handedness is associated with less common input to spinal motor neurons innervating different hand muscles. Journal of Neurophysiology, 128(4), 778–789.

49. Myers, L. J., Erim, Z., & Lowery, M. M. (2004). Time and frequency domain methods for quantifying common modulation of motor unit firing patterns. Journal of NeuroEngineering and Rehabilitation, 1, 1–12.

50. Napier, J. R. (1956). The prehensile movements of the human hand. The Journal of Bone & Joint Surgery British Volume, 38(4), 902–913.

51. Negro, F., Holobar, A., & Farina, D. (2009). Fluctuations in isometric muscle force can be described by one linear projection of low-frequency components of motor unit discharge rates. The Journal of Physiology, 587, 5925–5938.

52. Negro, F., & Farina, D. (2012). Factors influencing the estimates of correlation between motor unit activities in humans. Frontiers in Neuroscience.

53. Negro, F., Keenan, K., & Farina, D. (2015). Power spectrum of the rectified EMG: when and why is rectification beneficial for identifying neural connectivity?. Journal of neural engineering, 12(3), 036008.

54. Negro, F., Muceli, S., Castronovo, A. M., Holobar, A., & Farina, D. (2016). Multi-channel intramuscular and surface EMG decomposition by convolutive blind source separation. Journal of Neural Engineering, 13(2), 026027.

55. Negro, F., Yavuz, U. Ş., & Farina, D. (2016). The human motor neuron pools receive a dominant slow-varying common synaptic input. The Journal of physiology, 594(19), 5491–5505.

56. Negro, F., Laine, C. M., Mayer, F., Martinez-Valdes, E., Falla, D., & Farina, D. (2017). Tracking motor units longitudinally across experimental sessions with high-density surface electromyography.

57. Oliveira, A. S., & Negro, F. (2021). Neural control of matched motor units during muscle shortening and lengthening at increasing velocities. Journal of Applied Physiology, 130(6),

58. Ó’Reilly, D., & Delis, I. (2022). A network information theoretic framework to characterise muscle synergies in space and time. Journal of Neural Engineering, 19(1), 016031.1798–1813. 10.1152/japplphysiol.00043.2021

59. O’Reilly, D., & Delis, I. (2024). Dissecting muscle synergies in the task space. Elife, 12, RP87651.

60. Santello, M., Bianchi, M., Gabiccini, M., Ricciardi, E., Salvietti, G., Prattichizzo, D., … & Bicchi, A. (2016). Hand synergies: Integration of robotics and neuroscience for understanding the control of biological and artificial hands. Physics of life reviews, 17, 1–23.

61. Schieber, M. H. (1990). How might the motor cortex individuate movements?. Trends in neurosciences, 13(11), 440–445.

62. Schieber, M. H., & Poliakov, A. V. (1998). Partial inactivation of the primary motor cortex hand area: effects on individuated finger movements. Journal of Neuroscience, 18(21), 9038–9054.

63. Simpson, T. G., Godfrey, W., Torrecillos, F., He, S., Herz, D. M., Oswal, A., … & Tan, H. (2024). Cortical beta oscillations help synchronise muscles during static posture holding in healthy motor control. NeuroImage, 298, 120774.

64. Tinazzi, M., Farina, S., Tamburin, S., Facchini, S., Fiaschi, A., Restivo, D., & Berardelli, A. (2003). Task-dependent modulation of excitatory and inhibitory functions within the human primary motor cortex. Experimental brain research, 150(2), 222–229.

65. Tower, S. S. (1940). Pyramidal lesion in the monkey. Brain, 63(1), 36–90.

66. Tuthill, J. C., & Azim, E. (2018). Proprioception. Current Biology, 28(5), R194–R203.

67. Valero-Cuevas, F. J., Johanson, M. E., & Towles, J. D. (2003). Towards a realistic biomechanical model of the thumb: the choice of kinematic description may be more critical than the solution method or the variability/uncertainty of musculoskeletal parameters. Journal of biomechanics, 36(7), 1019–1030.

68. Watanabe, R. N., & Kohn, A. F. (2015). Fast oscillatory commands from the motor cortex can be decoded by the spinal cord for force control. Journal of Neuroscience, 35(40), 13687–13697.

69. Wu, C. C., Lin, Y. T., Chen, Y., Chen, Y. C., & Hwang, I. S. (2025). Blood flow restriction modulates common drive to motor units and force precision: Implications for neuromuscular coordination. European Journal of Applied Physiology, 1–12.

70. Yang, Y., Solis-Escalante, T., Van de Ruit, M., Van der Helm, F. C., & Schouten, A. C. (2016). Nonlinear coupling between cortical oscillations and muscle activity during isotonic wrist flexion. Frontiers in computational neuroscience, 10, 126.

71. Yao, B., Klein, C. S., Hu, H., Li, S., & Zhou, P. (2018). Motor unit properties of the first dorsal interosseous in chronic stroke subjects: Concentric needle and single fiber EMG analysis. Frontiers in Physiology, 9, 1587.

